# Speckle contrast optical spectroscopy improves cuffless blood pressure estimation compared to photoplethysmography

**DOI:** 10.1101/2024.08.16.608163

**Authors:** Ariane Garrett, Byungchan Kim, Nil Z. Gurel, Edbert J. Sie, Benjamin K. Wilson, Francesco Marsili, John P. Forman, Naomi M. Hamburg, David A. Boas, Darren Roblyer

## Abstract

Continuous and non-invasive blood pressure (BP) monitoring has the potential to greatly improve hypertension diagnosis and management, along with enabling valuable personal health monitoring in the population at large. Existing methods rely on cuff-based sphygmomanometers, which are cumbersome and disrupt sleep. Despite extensive research utilizing photoplethysmography (PPG), which is integrated into many consumer wearables, errors in continuous BP estimation are generally considered unacceptable for general use. We developed a high-speed (390 Hz) Speckle Contrast Optical Spectroscopy (SCOS) system to measure the cardiac blood flow waveform simultaneously with the PPG signal at high temporal resolution. The system utilized high speed multiplexed detection of optical speckle patterns on the wrist and finger, enabling the extraction of novel features related to BP. In comparison to PPG alone, SCOS demonstrated a notable 31% improvement (p = 3.45 * 10^-7^) in systolic BP estimation when integrated into subject-specific machine-learning models. The resulting errors were remarkably low (systolic BP: 0.06+/- 2.88 mmHg, diastolic BP: 0.09 +/-2.14 mmHg) across a wide range of BP variations (range SBP: 89–284 mmHg). Importantly, this improvement was sustained several weeks later within a re-measured cohort, indicating highly robust BP predictions. Looking ahead, the use of SCOS for blood flow measurements holds the potential to substantially enhance BP estimations compared to conventional PPG-based methods.

## Introduction

Hypertension is the leading cause of cardiovascular disease (CVD) and affects nearly half of adults in the Unites States (1). The current gold standard for noninvasive BP estimation is the cuff-based sphygmomanometer, which can only provide intermittent measures of BP (2). This prohibits continuous BP monitoring, making it difficult to measure daily BP fluctuations. Frequent BP measurements outside of the clinic provide a more robust assessment of BP than single measurements acquired in the clinician’s office (3) and the difference between daytime and nighttime BP is strongly correlated with cardiovascular risk prediction (4–6). However, current cuff-based BP measurements are disruptive to patient’s daily lives, limiting frequent measurements outside of the clinic. In the general population, continuous BP monitoring has the potential to identify those at risk, potentially leading to earlier and more effective interventions.

Much of the work towards noninvasive continuous BP monitoring has centered around estimating BP using the photoplethysmography (PPG) signal. The PPG signal arises from changes in hemoglobin absorption during each cardiac pulse, providing a relative measure of blood volume changes (7). PPG has traditionally been utilized for heart rate monitoring, and when conducted with at least two wavelengths, can provide pulse oximetry measurements. More recently, PPG signals have been integrated into consumer smart watches, enabling continuous heart rate and pulse oximetry measurements outside of the clinical setting (8). The ubiquitous nature of the PPG signal in both the clinic and daily life makes it an ideal candidate for continuous, noninvasive BP estimation. There has been much effort towards this goal, and various strategies have been applied. Physiological models such as the Windkessel and the Moens-Korteweg models have used the PPG signal as an input to estimate BP (9,10). Alternatively, features extracted from the PPG waveform have been shown to correlate with BP (11–16). These features have been used as inputs to a wide range of machine learning techniques for BP estimation (16–21).

Other techniques such as tonometry and ultrasound have also been explored for cuff-less blood pressure monitoring (22–26). Ultrasound uses ultrasonic waves to generate images of arteries, then generates a pulse waveform based on the change in diameter of the arteries with each heartbeat. Provided there is a BP calibration point, blood pressure can then be calculated using physiological models (27). Ultrasound can penetrate more deeply into tissue than optical techniques, allowing direct measurements of major arteries. Tonometry measurements of blood pressure require applanation of the target artery (typically radial) and use a transducer to measure the intra-arterial pressure wave (28). Compared to PPG, both ultrasound and tonometry are more sensitive to motion and require precise placement of the probe above the target artery (24,29) .

While substantial progress has been made using noninvasive BP techniques, errors remain unacceptable for clinical adoption (26,30,31). Additionally, each of these noninvasive BP techniques provides a single waveform related to either blood volume, vessel diameter, or pulse pressure. Taken individually, these waveforms are likely limited in their ability to fully capture the cardiovascular state. New multi-modal measurements may be needed to improve BP predictions over what is currently available.

In this work we investigated whether the simultaneous measurement of blood flow and blood volume in the periphery (i.e. finger and wrist) could improve on cuffless BP estimates. While the PPG signal is related to changes in pulsatile blood volume in tissue, it is insensitive to the changes in blood flow velocity that occurs during the cardiac cycle. It has long been known that both blood flow and volume changes better characterize the cardiovascular state (32), but non-invasive blood flow measurements capable of resolving blood flow changes within the cardiac cycle have been limited. To this end, we developed a new high-speed version of SCOS to measure blood flow changes at the cardiac rate. SCOS captures the dynamic interference patterns, or speckle patterns, arising from the scattering of laser light on tissue, which are directly influenced by the blood flow (33–35). By acquiring speckle images at a sufficiently high sampling rate, SCOS reveals the pulsatile blood flow index (BFi) waveform (36,37). The PPG and BFi waveforms are distinct (37), providing opportunities for new features that relate to cardiovascular health and BP (38). Furthermore, the BFi waveform is more resilient to noise during conditions of low tissue perfusion compared to the PPG signal (37), and is less sensitive to temperature and motion (39). All of this suggests that the BFi waveform measured using SCOS contains abundant, yet largely unexplored features that are relevant to BP estimation.

Here we describe a new custom high-speed (390 Hz) SCOS system that was used to acquire measurements at both visible (532 nm) and near infrared (NIR, 808 nm) wavelengths from 30 subjects before, during and after a leg press exercise designed to induce changes in BP. The high temporal resolution of the collected waveforms provides a variety of shape, time, and frequency-domain features that were used in subject-specific machine learning models to compute continuous BP. The additional information provided by SCOS improved BP estimates substantially compared to PPG alone in both leg press and cold pressor exercises. The patient-specific models continued to provide highly accurate BP predictions even after remeasurement approximately 20 weeks later. The addition of blood flow information from high-speed SCOS provides a new avenue for highly accurate, non-invasive, cuffless, and continuous BP measurements.

## Results

### Theory of Operation

SCOS utilizes long coherence laser illumination at wavelengths sensitive to red blood cells scattering and hemoglobin absorption to measure changes in blood flow and volume. During diastole, blood volume in the peripheral tissue compartments is at the lowest point of the cardiac cycle, resulting in decreased absorption from hemoglobin and increased measured intensity on the detector (Figure 1A). In addition, the blood flow is at a minimum, reducing the rate of speckle fluctuations measured by the detector. In practice, the images are acquired at a specific exposure time (in our case, 2.5ms). The rate of speckle fluctuations can therefore be quantified by measuring the blurriness of the speckle pattern, or speckle contrast (K). During diastole, the rate of speckle fluctuations is lower, so the blurriness is decreased and speckle contrast is increased. Conversely, during systole, blood volume is maximal, resulting in increased hemoglobin and decreased measured intensity. Blood flow is also increased, resulting in increased speckle fluctuations, increased blurriness of the speckle pattern, and decreased speckle contrast (Figure 1A). To quantify changes in blood flow, speckle contrast is calculated as the standard deviation of the speckle image divided by the mean intensity of the image. Sources of variance unrelated to blood flow changes are subtracted from the speckle contrast, and the relative blood flow index is calculated as 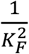 (Figure 1B). The blood volume (PPG) signal is calculated using the intensity of the speckle image, 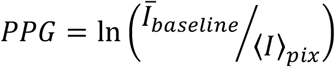 (Figure 1B). Details about the imaging processing are included in the methods section. When speckle images are acquired at a sufficient sampling rate, the changes in blood flow and volume throughout the entire cardiac pulse can be resolved (Figure 1C). The blood flow (BFi) and volume (PPG) signals originate from different physiological sources and thus exhibit different morphology and timing characteristics (Figure 1C, 1F).

**Figure 1.**
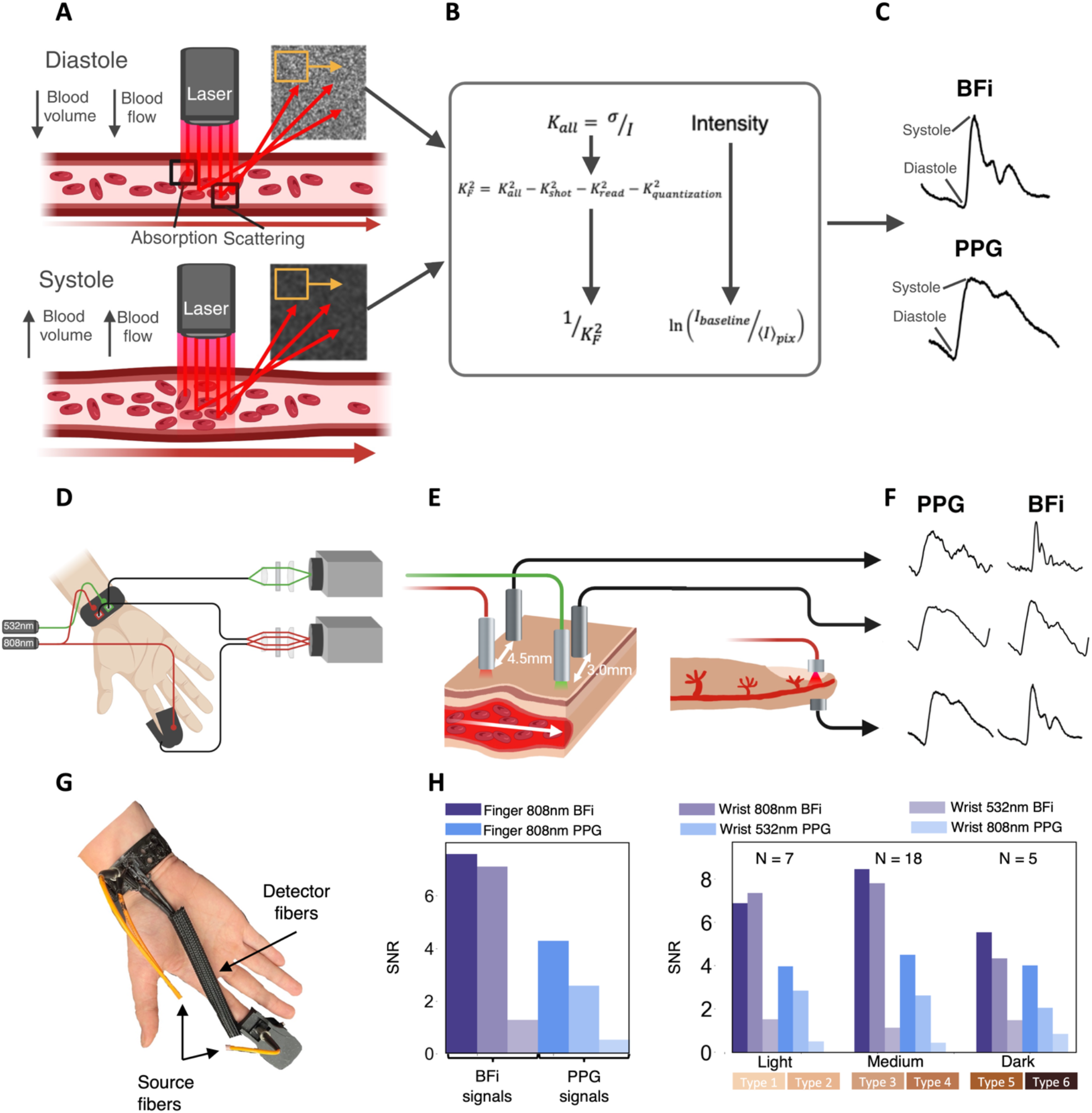
Device design. **a**. Principle of SCOS measurement. As blood flow and volume increase during systole, speckle contrast and intensity decrease. **b**. Data processing pipeline. **c**. Representative BFi and PPG waveforms during one cardiac pulse. **d**. Measurement set up. **e**. 532 nm and 808 nm reflective measurements are acquired at the wrist. An 808 nm transmission measurement is acquired at the finger. **f**. PPG and BFi waveforms at each measurement location. **g**. Wrist and finger SCOS devices placed on a representative subject. **h**. SNR comparison between BFi and PPG signals at each measurement location and for subjects of different Fitzpatrick numbers.

### Device design

The optical configuration (Figure 1D) comprised two long coherence laser sources, operating at wavelengths of 532 nm (Crystalaser CL532-075-SO) and 808 nm (Crystalaser DL808-100-S30), both coupled into multimode fibers (Thorlabs M42L01) with a numerical aperture of 0.22 and a core diameter of 50 μm. The two wavelengths chosen target distinct tissue depths. The 532 nm light is more significantly absorbed by tissue and penetrates less deeply, and has been commonly employed in wrist PPG measurements (40). In contrast, the 808 nm laser penetrates deeper into the tissue and is the isosbestic point between oxy- and deoxy-hemoglobin absorption, enabling monitoring of variations in total blood volume irrespective of the individual changes in oxy- and deoxy-hemoglobin concentrations. To illuminate both the finger and wrist with 808 nm light, a 50:50 fiber optic splitter was used, with one of the source fibers placed on the finger and the other on the wrist. The 532 nm source fiber was positioned 1 cm apart from the 808 nm source on the wrist. We collected the scattered light using three detector fibers, each corresponding to a source fiber. For finger measurements, the detector was placed on the opposite side of the finger relative to the source fiber to measure light transmission through the finger. For the wrist measurements, the detector fiber was situated at distances of 3.0 mm and 4.5 mm from the source for the 532 nm and 808 nm wrist measurements, respectively. The three detector fiber bundles have 3770 individual fibers per bundle, with a fiber core diameter of 37 μm and a numerical aperture of 0.66 (Fiber Optics Technology, Inc, FTIMG25911). These detection fibers were imaged onto two cameras (Basler Boost) using two plano-convex lenses (Edmund Optics, #67-581 and #67-531). Camera 1 captured the 808 nm light, while Camera 2 recorded the 532 nm light. The frame rate of both cameras was 390 Hz. Between the two plano-convex lenses prior to Camera 2, a short-pass filter with a cutoff wavelength of 700 nm (Thorlabs FELH0700) was placed to block 808 nm light. Similarly, a long-pass filter with a cutoff wavelength of 700 nm (Thorlabs FESH0700) was positioned before Camera 1 to prevent 532 nm light from entering (Figure 1D). Although PPG and BFi pulse waveforms (PWFs) can be obtained from all three measurement locations, the BFi PWFs from the 532 nm wrist measurement and the PPG PWFs from the 808 nm wrist measurement were too noisy in the majority of subjects to reliably extract features. Therefore, all data analysis in this paper was performed using the PPG and BFi PWFs from the finger, the PPG signal from the 532 nm wrist measurement, and the BFi signal from the 808 nm wrist measurement.

Figure 1F shows sample BFi and PPG waveforms from each measurement location. The BFi waveforms have higher signal to noise ratio (SNR) than the PPG waveforms at each measurement location and across different skin tones (Figure 1H). Example time traces showing the differences in noise between finger BFi and PPG signals are shown in Supplementary Figure 8 to further highlight the improved SNR of the BFi waveforms.

### Measurement protocol

SCOS measurements were acquired from 30 healthy subjects. Demographic information is shown in Supplementary Figure 1. Each optical measurement consisted of 532 nm and 808 nm reflective SCOS measurements on the wrist, and an 808 nm transmissive measurement on the finger. BP was simultaneously collected from the contralateral arm using the Finapres Nova (Finapres Medical Systems) continuous BP monitor (41,42). The measurement protocol is shown in Figure 2A. Briefly, after the leg press weight was identified, an initial one-minute baseline measurement was taken. Then, subjects completed a four-minute isometric leg press exercise using the previously selected weight while BP and optical measurements were obtained simultaneously. Subjects then rested for 15 minutes and a final one-minute recovery measurement was obtained. For five subjects, an additional cold pressor exercise was used to induce BP changes. The same procedure was followed as the leg press protocol, but instead of performing a leg press the subjects were asked to submerge their left hand in a bucket of ice water for two minutes or as long as was tolerable.

**Figure 2.**
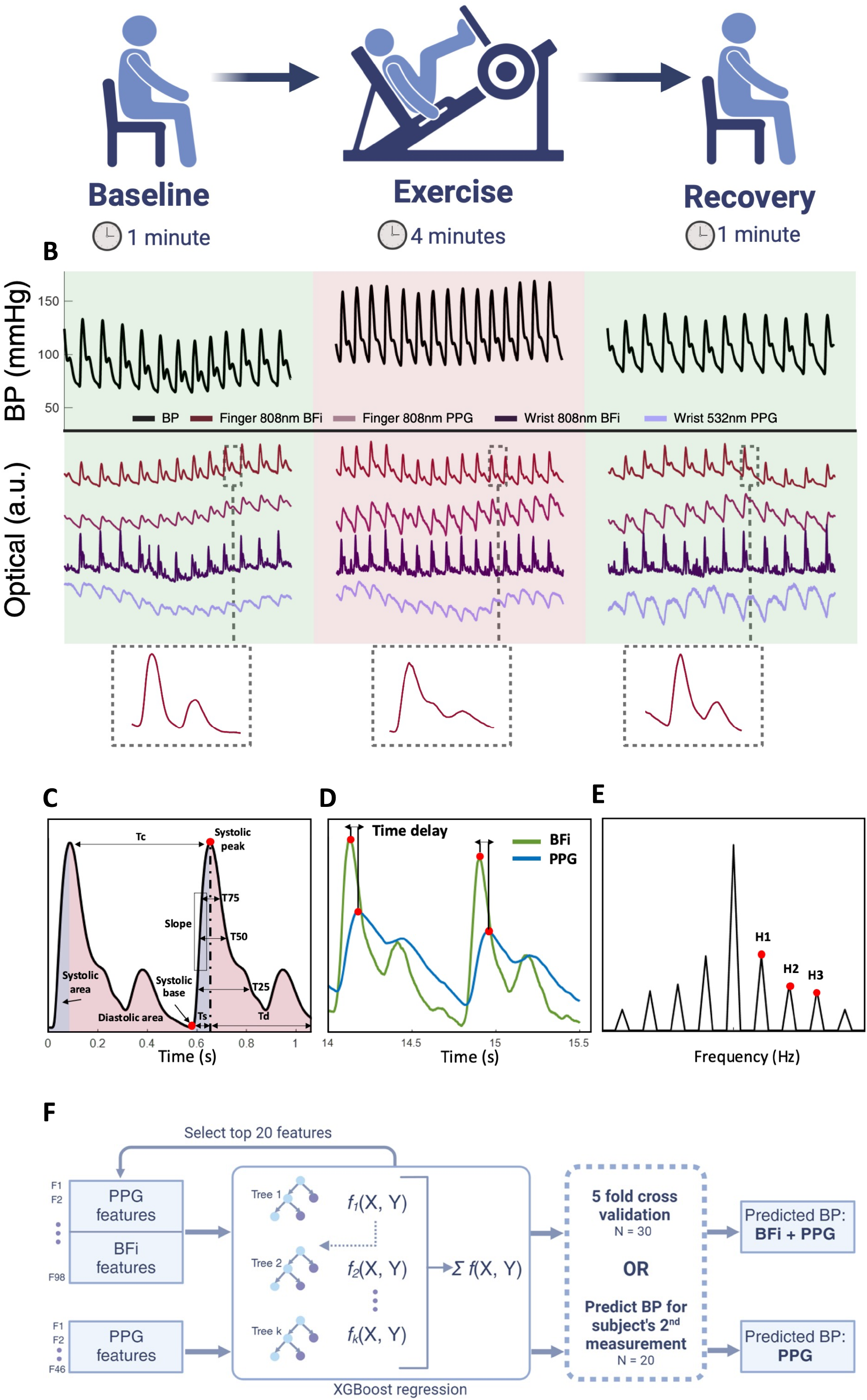
Measurement protocol and data analysis. **a**. Leg press measurement protocol. **b**. BP waveforms obtained from the Finapres and optical waveforms from the SCOS device during baseline, exercise, and recovery. Individual finger 808 nm BFi waveforms highlighting the morphology changes between baseline, exercise, and recovery are shown below the optical waveforms. **c**. Time and area related features shown on a representative BFi waveform Tc: length of cardiac cycle. Td: Diastolic time. Ts: Systolic time. T75, T50, T25: width of pulse at 75%, 50%, and 25% of the pulse amplitude, respectively. **d**. BFi and PPG waveforms from the finger highlighting the time delay between the systolic peaks of BFi and PPG waveforms, which is used as a feature. **e**. Frequency domain features. H1, H2, H3 are harmonic peaks. **f**. BP estimation model.

### BP estimation model

26 unique features were extracted for each measurement type (BFi and PPG), along with 14 joint features combining information between BFi and PPG (Supplementary Table 2). These features are illustrated in Figure 2C-E. A quality control algorithm was used to remove pulses that exceed a noise threshold from the analysis (described in Methods section). To estimate BP, a supervised machine learning model was used, eXtreme Gradient Boosting (XGBoost), which implements gradient boosted decision trees (Figure 2F). It has previously been used to estimate BP from PPG signals and has improved calculation speed and generalizability compared with other machine learning models (43).

### BP estimation using five-fold cross validation

First, BP was estimated for the original 30 subjects. Two XGBoost models were trained. For the first model, only PPG features were used as inputs (henceforth referred to as PPG model), and for the second model, both BFi and PPG features were used as inputs (henceforth referred to as BFi + PPG model) (Figure 2F). In total, there are 118 features input to the BFi + PPG model, and 52 features input to the PPG model. XGBoost’s built-in feature selection was used to select the top features for each model to ensure each model was trained on the same number of features and to reduce overfitting. To select the optimal number of features, errors were calculated for PPG and BFi + PPG models trained on feature numbers ranging from 1-40 (Figure 3B). The BFi + PPG model consistently outperformed the PPG model regardless of the number of features used in the model. We chose to use a maximum of 25 features, to balance minimizing the error with overfitting. For each subject, the data for that subject’s measurement was shuffled and BP was predicted using 5-fold cross validation (Figure 3A). The output BP was averaged to estimate a BP every five pulses (roughly every 5 seconds, depending on subject heart rate). The same procedure was followed for the five subjects measured during a cold pressor perturbation.

**Figure 3.**
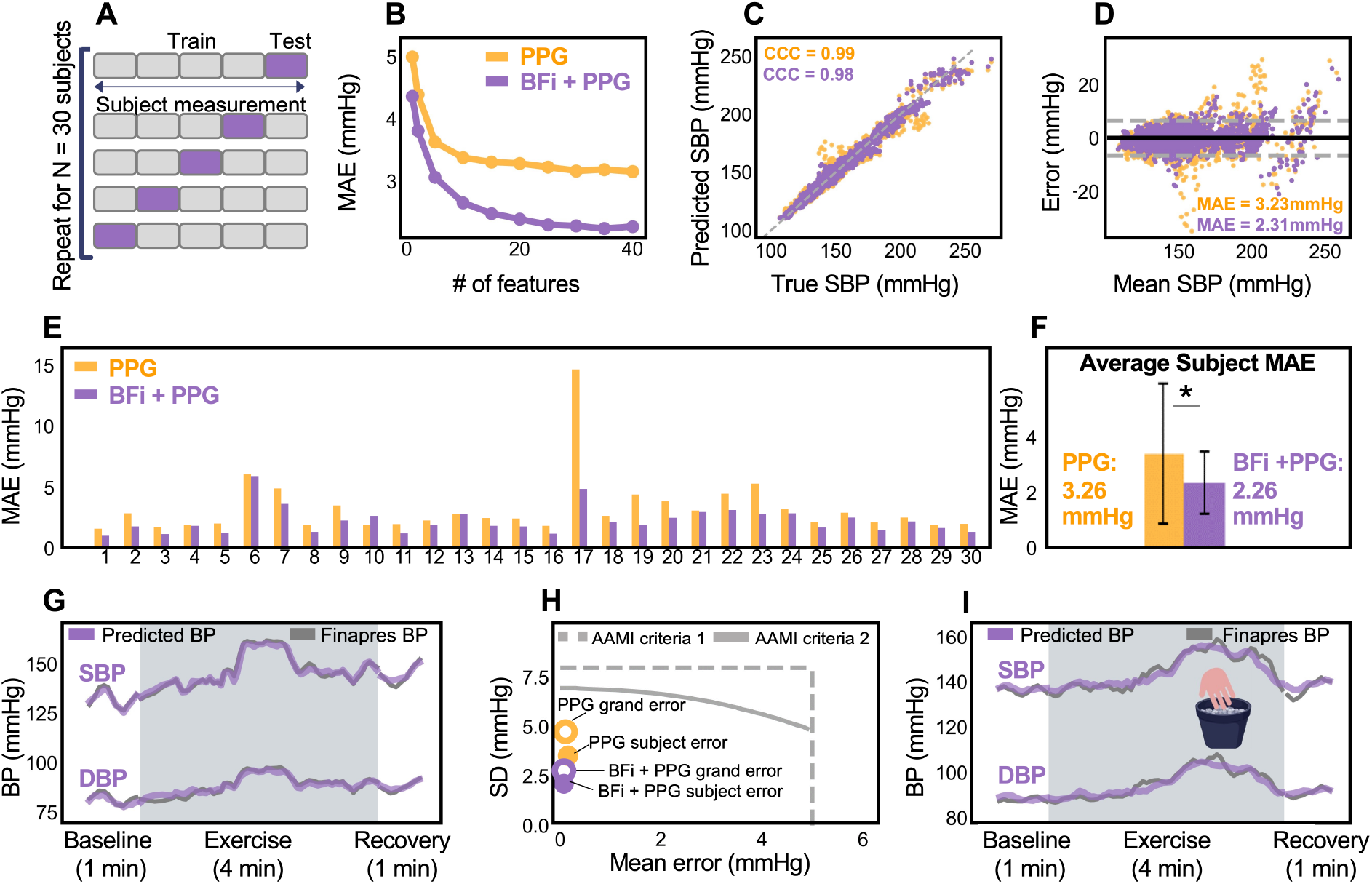
Results from 5-fold cross validation BP measurement. **a**. Subject’s BP is predicted using five-fold cross validation. **b**. The BFi + PPG model consistently predicts SBP with a lower MAE than the PPG model, regardless of the number of features used in the model. A maximum of 25 features was selected. **c**. Correlation between predicted and true SBP for the PPG model and BFi + PPG model (overlayed). CCC: concordance correlation coefficient. **d**. Bland – Altman plot for the PPG model and BFi + PPG model (overlayed) showing grand errors. Mean and standard deviations lines are for the BFi + PPG model. MAE: mean absolute error. ME: mean error. **e**. MAE for all 30 subjects showing improvement in MAE with the BFi + PPG model in the majority of subjects. Subject MAE are shown in plot. **f**. Average subject MAE for PPG and BFi + PPG models. The BFi + PPG model has a significantly lower MAE. **g**. Time trace for subject 1 showing predicted SBP and DBP during baseline, the leg press exercise, and recovery. **h**. PPG and BFi + PPG grand and subject SBP errors shown in context of AAMI criteria 1 and 2. **i**. Time trace for subject 1 showing predicted SBP and DBP during baseline, the cold pressor exercise, and recovery.

BP errors can be expressed as both a mean of the errors from each individual pulse (grand errors) and as a mean of the errors from each individual subject (subject errors). Grand errors weight the errors from each pulse equally, whereas subject errors weight the errors from each subject equally. The Association for the Advancement of Medical Instrumentation/European Society of Hypertension/International Organization for Standardization (AAMI/EHS/ISO ) guidelines specify different criteria for each type of error, so results from both are shown in Figures 3 and 4. The PPG model estimated systolic BP (SBP) with a grand mean absolute error (MAE) of 3.23 mmHg (Figure 3D), whereas the BFi + PPG model predicted SBP with a MAE of 2.31 mmHg (Figure 3E), a 29% improvement in MAE compared to the PPG-only model. The PPG model estimated systolic BP (SBP) for each subject with a mean absolute error (MAE) of 3.26 mmHg (Figure 3E-F), whereas the BFi + PPG model predicted SBP with a MAE of 2.26 mmHg (Figure 3E-F), a 31% improvement in MAE compared to the PPG-only model (p = 3.45 * 10^-7^). The PPG-only model estimated cold pressor SBP values with a MAE of 3.16 mmHg, whereas the BFi + PPG model predicted cold pressor SBP with a MAE of 2.33 mmHg, a 26% improvement in MAE compared to the PPG-only model. Both SBP and DBP results from the PPG and BFi + PPG models met the AAMI/EHS/ISO criterion.

**Figure 4.**
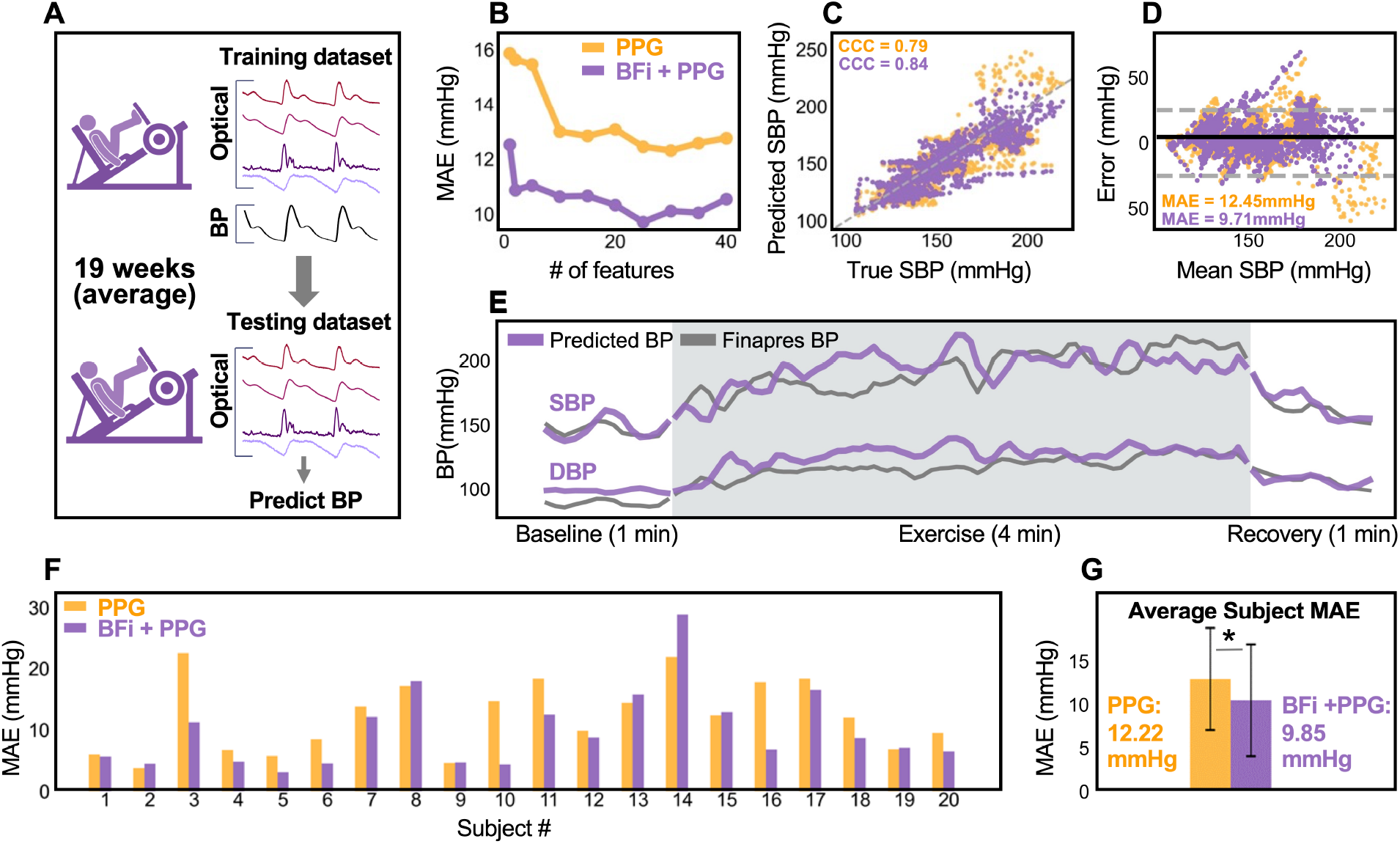
Results from longitudinal BP measurement. **a**. Subject’s BP is predicted using models trained on the subjects first measurement. **b**. The BFi + PPG model consistently predicts SBP with a lower MAE than the PPG model, regardless of the number of features used in the model. A maximum of 25 features was selected. **c**. Correlation between predicted and true SBP for the PPG model and BFi + PPG model (overlayed). CCC: concordance correlation coefficient. **d**. Bland – Altman plot for the PPG model and BFi + PPG model (overlayed) showing grand errors. Mean and standard deviations lines are for the BFi + PPG model. MAE: mean absolute error. ME: mean error. **e**. Time trace for subject 7 showing predicted SBP and DBP during baseline, the leg press exercise, and recovery **f**. MAE for all 20 subjects showing improvement in MAE with the BFi + PPG model in the majority of subjects. Subject MAE are shown in plot. **g**. Average subject MAE for PPG and BFi + PPG models. The BFi + PPG model has a significantly lower MAE.

Cold pressor results are shown in Supplementary Figure 4. Diastolic blood pressure (DBP) results are shown in Supplementary Figure 5.

### Longitudinal BP estimation

Next, BP was estimated for the 20 subjects who were remeasured an average of 18.8+/- 11.3 weeks following the first measurement session. Each subject was remeasured following the same leg press protocol described in Figure 2A. As with the original 30 subjects, both a BFi+PPG model and a PPG-only model were used to estimate BP and feature selection was used to select the top 25 features for each model. However, in this case the XGBoost models were trained on the subject’s first measurement session and used to predict BP for the second measurement session. This is more challenging because the measurements take place several weeks apart and changes in probe placement and subject physiology could impact the ability of the model to accurately predict BP. To mitigate these effects, output BP was calibrated using the 1^st^ and last BP measurements from the second measurement. Results are shown in Figure 4. Although estimation errors increased, the BFi + PPG model still outperformed the PPG model. The SBP grand MAE for the PPG only model was 12.45 mmHg, whereas the SBP grand MAE for the BFi + PPG model was 9.71 mmHg, a 22% improvement in MAE compared to the PPG-only model. The SBP subject MAE for the PPG only model was 12.22 mmHg, whereas the SBP grand MAE for the BFi + PPG model was 9.85 mmHg, a 19% improvement in MAE compared to the PPG-only model (p = 0.01). The DBP results are shown in Supplementary Figure 6.

## Discussion

Existing non-invasive BP devices have used a variety of physiological waveforms, including the PPG, ultrasound, or tonometry waveforms, which measure blood volume changes, vessel diameter, or pulse pressure respectively. Since BP is influenced by a variety of cardiovascular parameters, measurement from single modalities may limit the achievable accuracy of current methods, which has slowed adoption. To our knowledge, only one PPG based BP monitor has been approved by the FDA (44,45), and rigorous efforts to assess FDA cleared wearable cuffless BP devices have found that they frequently are not able to provide greater BP measurement accuracy than a baseline model, which only uses the calibration and time of day (26,31). Multi-modal techniques may provide a richer picture of the cardiovascular state, and techniques that have combined, for example, PPG and electrocardiogram (ECG) waveforms, have shown promising results (2,26,46). Here we pursued a new strategy in which blood flow information, enabled by high-speed SCOS, was combined with the PPG waveform to enable highly accurate BP estimates. The blood flow (BFi) and PPG measurements are captured simultaneously with the SCOS system, providing an inherently multi-modal platform.

SCOS allows the extraction of BFi waveforms which are dependent on blood flow changes, and are morphologically and temporally distinct from the PPG waveform (37). It has been previously shown that the BFi waveform exhibits correlations with various cardiovascular health parameters and age, and is more resilient in terms of signal to noise ratio (SNR) during conditions of low tissue perfusion compared to the PPG signal (37). It has also been shown that BFi is superior to PPG in measurements of heart rate variability (HRV), possibly due to its reduced sensitivity to temperature and motion artifacts (39). Here we observed that the BFi waveforms had higher SNR than the PPG waveforms across different skin tones (Figure 1H). That being said, we also observed that the 532 nm BFi signals from the wrist and the 808 nm PPG signals from the wrist were noisy in the majority of subjects. It has previously been hypothesized that the 532 nm PPG signal primarily arises from compression of the capillary beds by underlying arteries, rather than changes in blood volume within the capillaries themselves (40). In this case, the blood flow signal through the capillaries may be minimal although the blood volume signal remains strong, contributing to a noisy BFi and robust PPG signal. In the 808 nm reflectance PPG measurement on the wrist, clear inverted PPG waveforms were observed in some subjects, although the BFi signals extracted from the same dataset were not inverted. We hypothesize that the presence of inverted signals in the 808 nm PPG reflectance measurements may have led to increased noise. Further investigation is needed to confirm these hypotheses. Overall, the increased SNR of BFi signals offers a significant advantage over PPG and could facilitate more accurate BP estimation over a wide range of skin tones and perfusion conditions.

Novel features extracted from SCOS measurements increase the depth of information available for the assessment of the cardiovascular state. Prior work showed that the analysis of the pulsatile speckle waveform enhanced the classification of the progression of peripheral artery disease compared to traditional assessment methods (47). In our own prior work we demonstrated that individual features combining both BFi and PPG information were more strongly correlated with BP than PPG features alone (38). In this work we estimated a wide range of BP values (min SBP 89, max 284 mmHg) using two different methods, first by shuffling an individual subject’s measurement and performing 5-fold cross validation (48). Under this prediction strategy there was a substantial 29% improvement in SBP grand MAE and a 31% improvement in subject MAE (p = 3.45 * 10^-7^) when both BFi and PPG features were input. We also estimated BP in a more challenging framework in which subjects were remeasured several weeks after their first measurement. While this type of repeat measurement is not common in prior literature, with many studies utilizing single measurement sessions (20,49–53), it is a more clinically relevant prediction framework. Increases in BP estimation errors were anticipated because probe placement and physiological changes between the first and second measurement likely affect model accuracy. For the remeasured subjects we found that although overall errors increased, the BFi + PPG model improved grand MAE by 22% compared with the PPG-only model, and improved subject MAE by 19% (p = 0.01). This demonstrated that the improvement in BP estimation using SCOS persisted under different testing frameworks. We also found that on average, BFi features were strongly favored by the prediction model and had greater importance compared to PPG features (Supplementary Figure 8). The BFi + PPG model also improved BP estimation during the cold pressor exercise. Compared to BP changes during exercise, which are driven by increases in cardiac output, BP changes during the cold pressor are due to vasoconstriction, (54,55). This suggests that the improvement in estimation error by BFi is consistent across different physiological BP perturbations.

### Outlook

Limitations of this study include a relatively small sample size and the inclusion of only normotensive subjects. Future work will aim to measure both normotensive and hypertensive subjects and expand the subject pool. In addition, miniaturizing the system into a wireless, wearable device would enable long term measurements during daily activities, such as sleep. BP errors may be further improved by the use of different machine learning models or deep learning models that were not explored within the scope of this study.

## Conclusion

High-speed SCOS provides a multi-modal measurement of peripheral blood volume and flow changes during the cardiac cycle. The combination of BFi and PPG waveforms led to substantial improvements in BP estimates compared to PPG alone. We went beyond standard practice by predicting BP during different perturbations and a longitudinal measurement and found that SCOS consistently improves BP estimation across different measurement conditions and prediction strategies. SCOS has the potential to greatly improve hypertension diagnosis and monitoring compared to current cuff-based sphygmomanometers, which cannot provide continuous BP monitoring.

## Methods

### Subject pool

A total of 30 healthy volunteers were recruited. The subject’s age, height, weight, and Fitzpatrick score were collected. The Fitzpatrick score was used to quantify skin tone (56). Each subject was asked to select a number between one and six depending on how easily their skin burns when exposed to sun, with 6 corresponding to never burning and the darkest skin tone, and 1 corresponding to always burning and the lightest skin tone. The mean +/- standard deviation (SD) age of the subjects was 32 +/- 15 years, and the mean Fitzpatrick number was 3.1 +/- 1.2. The mean SBP increase during the leg press exercise in the first measurement (N = 30 subjects) was 23.2 +/- 20.5 mmHg, and the mean DBP increase was 13.7 +/- 13.8 mmHg. The mean SBP increase during the cold pressor exercise (N = 5 subjects) was 22.7 +/- 15.2 mmHg and the mean DBP increase was 15.1 +/- 12.3 mmHg. The mean SBP increase during the leg press for the repeat measurement (N = 20 subjects) was 19.3 +/- 13.8 mmHg and the mean DBP increase was 11.6 +/- 10.1 mmHg. Distributions of SBP, DBP, and heart rate (HR) for each measurement are shown in Supplementary Figures 1-3.

### Data collection

All data was collected and processed at Boston University. The images were collected at 390 Hz and a 2.5 ms exposure time, and each image was 400 × 400 pixels. The Finapres measures BP at a sampling rate of 200 Hz, and this data was collected by a National Instruments DAQ (NI USB 6361) with an acquisition rate of 1000 Hz. The BP and optical data were then aligned in time using an exposure active transistor-transistor logic (TTL) signal from the camera. Prior to data collection, the wrist measurement location was selected by collecting images at a frame rate of 25 Hz while scanning the probe over the volar side of the wrist until cardiac pulses were visible. Then, subjects were asked to test various leg press weights. Based on the observed BP changes and subjects’ assessment of their physical capabilities, a weight was selected.

### Data processing

The data processing pipeline is shown in Figure 1B. To obtain the PPG time series, a 400 x 400 pixel region of interest (ROI) imaging the fiber bundle was selected on each camera. Intensity values in the ROI pixels and across frames were averaged to obtain a baseline intensity, 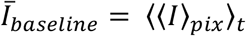, for the ROI. Then, the PPG signal was calculated as the natural log of the baseline intensity divided by spatially averaged intensity in each frame, 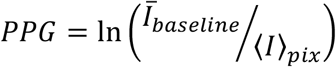. Optical blood flow measurements are calculated from squared speckle contrast, 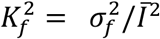, which is the spatial variance from the speckle patterns 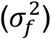 divided by squared spatial mean intensity 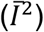. BFi is inversely proportional to the squared speckle contrast (57). However, spatial variance may be caused by other noise sources, including shot noise, read noise, spatial heterogeneity, and quantization noise (58). To reduce the effects of variance introduced by spatial heterogeneity in illumination, ROI was divided into 7x7 pixel windows with maximum overlap along both x and y axis. The raw squared contrast 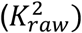 was calculated for each window by dividing each window’s spatial variance by squared mean intensity 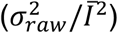. Other noise contributions were estimated (38) and subtracted from 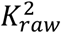 to isolate 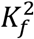 arising from the speckle patterns (Figure 1B). To obtain the BFi time series, speckle contrast and intensity were calculated for each image. For each window, spatial variance (*σ*^2^) and squared mean intensity were calculated to determine the measured squared contrast, *K*_*f*_. Then the BFi was calculated as 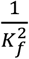 (Figure 1B).

### Peak identification

First, the individual peaks were identified by taking the first derivative of the time series and identifying the peaks in the differentiated signal using Matlab’s diff and findpeaks functions, respectively. The period of the cardiac cycle (Tc) was determined based on the heart rate, which was identified by the maximum of the Fourier transform of the signal within the range of 0.2 to 3.0 Hz. For each initially identified peak, the region 0.5*Tc before and 0.5*Tc after the identified peak was selected. This data represents an individual pulse waveform. This process was repeated to identify each of the individual pulse waveforms (PWFs), which were then averaged together to generate an average PWF.

### Quality control

To quantify the contribution of noise, individual cardiac waveforms were compared to the averaged waveform and PWFs that deviated significantly from the average waveform were removed. For this step, only the cardiac signal data 0.2 Tc before and 0.4 Tc after the systolic upslope at each pulse are considered. The average waveform is converted to a wavelet by subtracting the DC component and dividing by the amplitude of the waveform vector (square root of dot product of itself). For each individual cardiac waveform, the DC component is subtracted, then linearly fitted to the wavelet. To do so, the wavelet is scaled by the ratio between the dot product of the wavelet with the individual waveform and the dot product of the individual waveform with itself. The linearly fitted wavelet is subtracted from the original waveform to obtain the residual: *r* = *v* − *aw*; *a* = (*w* · *v*)/(*v* · *v*), where r = residual, a = ratio to scale wavelet by, w = trial averaged waveform’s wavelet, v = individual waveform. Then, the residual ratio is calculated as the residual’s power divided by the individual waveform’s power: *R* = (*r* · *r*)/(*v* · *v*), where R is the residual ratio. The residual ratio provides a value representing the proportion of noise power relative to the original individual waveform’s power. We use a threshold to classify individual waveforms, where the waveforms with residual ratio above the threshold are considered too noisy.

### Feature extraction

Many of the features are dependent on the identification of the base and maximum of the systolic peak. To identify the systolic peak, first we estimated approximate location of the peak on the individual pulse by finding the maximum systolic upslope from the first derivative of the signal and selecting the data 0.1 sec before and 0.2 sec after the max upslope. The systolic peak was then determined by Matlab’s findpeaks function applied over the selected data. The systolic base was defined as the intersecting tangent between two lines; the horizontal tangent at the minimum value 0.2 sec before the max upslope, and the tangent of the max upslope (59). Using the identified peak and trough values at each pulse, all features were determined except frequency domain features. To find the frequency domain features, the pulse waveform was padded and a Fourier transform was applied. Then, the peaks at the first, second, and third harmonics were identified and the ratios of these peaks were used as input features. Time points at which greater than 20% of features had been removed by the quality control were excluded from training and testing data. A table of all features is shown in Supplementary Table 2.

### Hyperparameter tuning

Hyperparameters were selected for each subject using Hyperopt, a distributed asynchronous hyperparameter optimization tool(60). Separate hyperparameters were selected for the PPG model and BFi + PPG model.

### Feature Selection

XGBoost’s built in feature selection tool was used to select the top 25 features. To analyze the top features chosen, feature importance for each subject’s top 25 features was summed across the 20 remeasured subjects. The top 25 features with the highest importance sum are shown in Supplementary table 3. To compare the importance of BFi and PPG features within individual subjects, the importances of features in each category were summed for each subject’s top 25 features.

Supplementary figure 8 shows the relative importances of BFi vs PPG features for each of the remeasured subjects.

### Statistical analysis

The Wilcoxon signed-rank test was used to calculate statistical significance in the difference between each subject’s mean absolute error in SBP prediction. Importantly, this test was only computed on the subject specific errors. Statistical tests were not performed on the grand (pulse) errors due to a lack of independence between individual pulse errors.

### SNR calculation

The SNR for each signal was calculated by first identifying each pulse as described above. The standard deviation of the peak values divided by the mean peak value was used to calculate the SNR.

### BP Calibration

Calibration was performed for the longitudinal BP estimation. The first and last predicted BP points were averaged and scaled to equal the average of the first and last Finapres BP points.

## References

1. Ostchega Y, Fryar CD, Nwankwo T, Nguyen DT. Hypertension Prevalence Among Adults Aged 18 and Over: United States, 2017-2018 Key findings Data from the National Health and Nutrition Examination Survey. 2017;

2. Sharma M, Barbosa K, Ho V, Griggs D, Ghirmai T, Krishnan SK, et al. Cuff-Less and Continuous Blood Pressure Monitoring: A Methodological Review. Vol. 5, Technologies. 2017.

3. Mancia G, Verdecchia P. Clinical value of ambulatory blood pressure: evidence and limits. Circ Res. 2015;116(6):1034–45.

4. Banegas JR, Ruilope LM, de la Sierra A, Vinyoles E, Gorostidi M, de la Cruz JJ, et al. Relationship between Clinic and Ambulatory Blood-Pressure Measurements and Mortality. N Engl J Med. 2018 Apr;378(16):1509–20.

5. Sherwood A, Hill LK, Blumenthal JA, Hinderliter AL. The Effects of Ambulatory Blood Pressure Monitoring on Sleep Quality in Men and Women With Hypertension: Dipper vs. Nondipper and Race Differences. Am J Hypertens. 2019 Jan;32(1):54–60.

6. Agarwal R, Light RP. The Effect of Measuring Ambulatory Blood Pressure on Nighttime Sleep and Daytime Activity—Implications for Dipping. Clin J Am Soc Nephrol. 2010;5(2):281.

7. Alian AA, Shelley KH. Photoplethysmography. Best Pract Res Clin Anaesthesiol. 2014;28(4):395– 406.

8. Elgendi M, Fletcher R, Liang Y, Howard N, Lovell NH, Abbott D, et al. The use of photoplethysmography for assessing hypertension. NPJ Digit Med. 2019;2(1):1–11.

9. Choudhury AD, Banerjee R, Sinha A, Kundu S. Estimating blood pressure using Windkessel model on photoplethysmogram. In: 2014 36th annual international conference of the IEEE engineering in medicine and biology society. IEEE; 2014. p. 4567–70.

10. Mukkamala R, Hahn JO, Inan OT, Mestha LK, Kim CS, Töreyin H, et al. Toward ubiquitous blood pressure monitoring via pulse transit time: theory and practice. IEEE Trans Biomed Eng. 2015;62(8):1879–901.

11. Block RC, Yavarimanesh M, Natarajan K, Carek A, Mousavi A, Chandrasekhar A, et al. Conventional pulse transit times as markers of blood pressure changes in humans. Sci Rep. 2020;10(1):1–9.

12. Elgendi M. Detection of c, d, and e waves in the acceleration photoplethysmogram. Comput Methods Programs Biomed. 2014;117(2):125–36.

13. Elgendi M, Liang Y, Ward R. Toward generating more diagnostic features from photoplethysmogram waveforms. Diseases. 2018;6(1):20.

14. Lin WH, Wang H, Samuel OW, Liu G, Huang Z, Li G. New photoplethysmogram indicators for improving cuffless and continuous blood pressure estimation accuracy. Physiol Meas. 2018;39(2):25005.

15. Addison PS. Slope transit time (STT): A pulse transit time proxy requiring only a single signal fiducial point. IEEE Trans Biomed Eng. 2016;63(11):2441–4.

16. Chowdhury MH, Shuzan MNI, Chowdhury MEH, Mahbub ZB, Uddin MM, Khandakar A, et al. Estimating blood pressure from the photoplethysmogram signal and demographic features using machine learning techniques. Sensors. 2020;20(11):3127.

17. Acciaroli G, Facchinetti A, Pillonetto G, Sparacino G. Non-Invasive Continuous-Time Blood Pressure Estimation from a Single Channel PPG Signal using Regularized ARX Models. In: 2018 40th Annual International Conference of the IEEE Engineering in Medicine and Biology Society (EMBC). EEE; 2018.

18. Duan K, Qian Z, Atef M, Wang G. A feature exploration methodology for learning based cuffless blood pressure measurement using photoplethysmography. In: 2016 38th Annual International Conference of the IEEE Engineering in Medicine and Biology Society (EMBC). IEEE; 2016.

19. Liang Y, Chen Z, Ward R, Elgendi M. Photoplethysmography and Deep Learning: Enhancing Hypertension Risk Stratification. Biosensors [Internet]. 2018;8(4):101. Available from: https://www.mdpi.com/2079-6374/8/4/101/pdf

20. Man PK, Cheung KL, Sangsiri N, Shek WJ, Wong KL, Chin JW, et al. Blood Pressure Measurement: From Cuff-Based to Contactless Monitoring. In: Healthcare. MDPI; 2022. p. 2113.

21. Qin C, Wang X, Xu G, Ma X. Advances in Cuffless Continuous Blood Pressure Monitoring Technology Based on PPG Signals. Biomed Res Int. 2022;2022.

22. Hosanee M, Chan G, Welykholowa K, Cooper R, Kyriacou PA, Zheng D, et al. Cuffless single-site photoplethysmography for blood pressure monitoring. J Clin Med. 2020;9(3):723.

23. Lin M, Zhang Z, Gao X, Bian Y, Wu RS, Park G, et al. A fully integrated wearable ultrasound system to monitor deep tissues in moving subjects. Nat Biotechnol. 2024;42(3):448–57.

24. Kenny JES. Wearable ultrasound for continuous deep-tissue monitoring. Nat Biotechnol. 2024;42(3):386–7.

25. Wang C, Li X, Hu H, Zhang L, Huang Z, Lin M, et al. Monitoring of the central blood pressure waveform via a conformal ultrasonic device. Nat Biomed Eng. 2018;2(9):687–95.

26. Mieloszyk R, Twede H, Lester J, Wander J, Basu S, Cohn G, et al. A Comparison of Wearable Tonometry, Photoplethysmography, and Electrocardiography for Cuffless Measurement of Blood Pressure in an Ambulatory Setting. IEEE J Biomed Heal informatics. 2022 Jul;26(7):2864–75.

27. Amado Rey AB. Estimation of Blood Pressure by Image-Free, Wearable Ultrasound. Artery Res. 2024;30(1):3.

28. Salvi P, Grillo A, Parati G. Noninvasive estimation of central blood pressure and analysis of pulse waves by applanation tonometry. Hypertens Res [Internet]. 2015;38(10):646–8. Available from: 10.1038/hr.2015.78

29. Deng M, Du C, Fang J, Xu C, Guo C, Huang J, et al. Flexible adaptive sensing tonometry for medical-grade multi-parametric hemodynamic monitoring. npj Flex Electron [Internet]. 2024;8(1):45. Available from: 10.1038/s41528-024-00329-9

30. Mukkamala R, Stergiou GS, Avolio AP. Cuffless blood pressure measurement. Annu Rev Biomed Eng. 2022;24:203–30.

31. Mukkamala R, Shroff SG, Landry C, Kyriakoulis KG, Avolio AP, Stergiou GS. The Microsoft Research Aurora Project: Important Findings on Cuffless Blood Pressure Measurement. Hypertension. 2023 Mar;80(3):534–40.

32. Westerhof N, Lankhaar JW, Westerhof BE. The arterial Windkessel. Med Biol Eng Comput [Internet]. 2009;47(2):131–41. Available from: https://link.springer.com/content/pdf/10.1007%2Fs11517-008-0359-2.pdf

33. Valdes CP, Varma HM, Kristoffersen AK, Dragojevic T, Culver JP, Durduran T. Speckle contrast optical spectroscopy, a non-invasive, diffuse optical method for measuring microvascular blood flow in tissue. Biomed Opt Express. 2014;5(8):2769–84.

34. Dragojević T, Hollmann JL, Tamborini D, Portaluppi D, Buttafava M, Culver JP, et al. Compact, multi-exposure speckle contrast optical spectroscopy (SCOS) device for measuring deep tissue blood flow. Biomed Opt Express. 2018;9(1):322–34.

35. Boas DA, Dunn AK. Laser speckle contrast imaging in biomedical optics. J Biomed Opt. 2010;15(1):11109.

36. Bi R, Du Y, Singh G, Ho JH, Zhang S, Attia ABE, et al. Fast pulsatile blood flow measurement in deep tissue through a multimode detection fiber. J Biomed Opt. 2020;25(5):55003.

37. Ghijsen M, Rice TB, Yang B, White SM, Tromberg BJ. Wearable speckle plethysmography (SPG) for characterizing microvascular flow and resistance. Biomed Opt Express. 2018;9(8):3937–52.

38. Garrett A, Kim B, Sie EJ, Gurel NZ, Marsili F, Boas DA, et al. Simultaneous photoplethysmography and blood flow measurements towards the estimation of blood pressure using speckle contrast optical spectroscopy. Biomed Opt Express. 2023;14(4):1594–607.

39. Dunn CE, Monroe DC, Crouzet C, Hicks JW, Choi B. Speckleplethysmographic (SPG) Estimation of Heart Rate Variability During an Orthostatic Challenge. Sci Rep. 2019;9(1).

40. Kamshilin AA, Margaryants NB. Origin of Photoplethysmographic Waveform at Green Light. Phys Procedia [Internet]. 2017;86:72–80. Available from: 10.1016/j.phpro.2017.01.024

41. Liu J, Yan BP, Zhang YT, Ding XR, Su P, Zhao N. Multi-wavelength photoplethysmography enabling continuous blood pressure measurement with compact wearable electronics. IEEE Trans Biomed Eng. 2018;66(6):1514–25.

42. Zadi AS, Alex RM, Zhang R, Watenpaugh DE, Behbehani K. Mathematical Modeling of Arterial Blood Pressure Using Photo-Plethysmography Signal in Breath-hold Maneuver. In: 2018 40th Annual International Conference of the IEEE Engineering in Medicine and Biology Society (EMBC). IEEE; 2018.

43. Chen T, He T, Benesty M, Khotilovich V, Tang Y, Cho H, et al. Xgboost: extreme gradient boosting. R Packag version 04-2. 2015;1(4):1–4.

44. Nachman D, Gepner Y, Goldstein N, Kabakov E, Ishay A Ben, Littman R, et al. Comparing blood pressure measurements between a photoplethysmography-based and a standard cuff-based manometry device. Sci Rep. 2020;10(1):16116.

45. BioBeat. U.S. Food and Drug Administration, Center for Drug Evaluation and Research. BioBeat Platform, BB-613WP Patch approval letter [Internet]. [cited 2023 May 12]. Available from: chrome-extension://efaidnbmnnnibpcajpcglclefindmkaj/https://www.accessdata.fda.gov/cdrh_docs/pdf21/K212153.pdf

46. Ghosh S, Banerjee A, Ray N, Wood PW, Boulanger P, Padwal R. Continuous blood pressure prediction from pulse transit time using ECG and PPG signals. In: 2016 IEEE Healthcare Innovation Point-Of-Care Technologies Conference (HI-POCT). 2016. p. 188–91.

47. Razavi MK, Flanigan DPT, White SM, Rice TB. A Real-Time Blood Flow Measurement Device for Patients with Peripheral Artery Disease. J Vasc Interv Radiol. 2021;32(3):453–8.

48. Sel K, Osman D, Huerta N, Edgar A, Pettigrew RI, Jafari R. Continuous cuffless blood pressure monitoring with a wearable ring bioimpedance device. npj Digit Med. 2023;6(1):59.

49. Saeed M, Villarroel M, Reisner AT, Clifford G, Lehman LW, Moody G, et al. Multiparameter Intelligent Monitoring in Intensive Care II (MIMIC-II): a public-access intensive care unit database. Crit Care Med. 2011;39(5):952.

50. Liang Y, Chen Z, Liu G, Elgendi M. A new, short-recorded photoplethysmogram dataset for blood pressure monitoring in China. Sci data. 2018;5(1):1–7.

51. Liu Q, Yan BP, Yu CM, Zhang YT, Poon CCY. Attenuation of Systolic Blood Pressure and Pulse Transit Time Hysteresis During Exercise and Recovery in Cardiovascular Patients. IEEE Trans Biomed Eng. 2014;61(2):346–52.

52. Schrumpf F, Frenzel P, Aust C, Osterhoff G, Fuchs M. Assessment of non-invasive blood pressure prediction from ppg and rppg signals using deep learning. Sensors. 2021;21(18):6022.

53. Kachuee M, Kiani MM, Mohammadzade H, Shabany M. Cuffless blood pressure estimation algorithms for continuous health-care monitoring. IEEE Trans Biomed Eng. 2016;64(4):859–69.

54. Mohammed LLM, Dhavale M, Abdelaal MK, Alam ABMN, Blazin T, Prajapati D, et al. Exercise-Induced Hypertension in Healthy Individuals and Athletes: Is it an Alarming Sign? Cureus. 2020 Dec;12(12):e11988.

55. Peckerman A, Hurwitz BE, Saab PG, Llabre MM, Mccabe PM, Schneiderman N. Stimulus dimensions of the cold pressor test and the associated patterns of cardiovascular response. Psychophysiology [Internet]. 1994;31(3):282–90. Available from: https://onlinelibrary.wiley.com/doi/abs/10.1111/j.1469-8986.1994.tb02217.x

56. Sachdeva S. Fitzpatrick skin typing: Applications in dermatology. Indian J Dermatol Venereol Leprol. 2009;75(1):93.

57. Fercher AF, Briers JD. Flow visualization by means of single-exposure speckle photography. Opt Commun. 1981;37(5):326–30.

58. Zilpelwar S, Cheng X, Sie E, Marsili F, Boas DA. Dynamic speckle model for simulating fiber-based laser speckle contrast imaging (fb-LSCI). In: Dynamics and Fluctuations in Biomedical Photonics XVIII. International Society for Optics and Photonics; 2021. p. 116410N.

59. Kazanavicius E, Gircys R, Vrubliauskas A, Lugin S. Mathematical methods for determining the foot point of the arterial pulse wave and evaluation of proposed methods. Inf Technol Control. 2005;34(1).

60. Bergstra J, Yamins D, Learning DCBTP of the 30th IC on M. Making a Science of Model Search: Hyperparameter Optimization in Hundreds of Dimensions for Vision Architectures [Internet]. Dasgupta S, McAllester D, editors. Vol. 28. PMLR; p. 115–23. Available from: http://proceedings.mlr.press/v28/bergstra13.pdf

