## Supplementary Materials for "Speckle contrast optical spectroscopy improves cuffless blood pressure estimation compared to photoplethysmography"

### Supplementary material

**Table 1 l Subject demographic information.** * indicates that the subject was remeasured for the longitudinal BP measurement. + indicates that the subject was measured for the cold pressor measurement. L* is lightness index measured from a colorimeter. BMI: body mass index

.

| Subject # | Age | Sex | Weight pressed (lbs) | Fitzpatrick # | L* | | BMI |
|---|---|---|---|---|---|---|---|
| | | | | | Finger | Wrist | |
| 1* | 28 | F | 110 | 3 | 57.15 | 63.04 | 25.5 |
| 2* | 27 | F | 90 | 2 | 56.36 | 67.06 | 19.3 |
| 3*+ | 24 | F | 110 | 3 | 55.94 | 62.77 | 23.2 |
| 4*+ | 25 | M | 140 | 2 | 54.31 | 63.74 | 23.1 |
| 5*+ | 25 | F | 100 | 4 | 51.58 | 67.23 | 23.3 |
| 6*+ | 40 | M | 140 | 3 | 56.94 | 63.71 | 27.3 |
| 7* | 40 | F | 130 | 4 | 58.86 | 65.69 | 26.6 |
| 8 | 25 | F | 130 | 3 | 57.68 | 67.64 | 21.6 |
| 9*+ | 25 | F | 90 | 3 | 52.7 | 63.22 | 21.0 |
| 10 | 24 | M | 180 | 2 | 50.2 | 61.41 | 27.0 |
| 11* | 23 | M | 160 | 6 | 56.13 | 57.34 | 22.3 |
| 12* | 24 | M | 170 | 3 | 54.81 | 61.94 | 22.4 |
| 13* | 22 | F | 90 | 4 | 54.57 | 58.79 | 21.2 |
| 14 | 18 | F | 110 | 1 | 56.69 | 69.38 | 26.9 |
| 15 | 79 | F | 70 | 3 | 51.93 | 52.87 | 22.7 |
| 16 | 23 | F | 140 | 2 | 57 | 64.91 | 24.8 |
| 17* | 27 | M | 200 | 3 | 55.42 | 60.52 | 23.1 |
| 18 | 46 | M | 150 | 4 | 52.71 | 54.56 | 21.3 |
| 19 | 25 | F | 150 | 3 | 58.97 | 65.69 | 19.0 |
| 20 | 36 | F | 120 | 3 | 52.19 | 59.25 | 21.9 |
| 21* | 67 | F | 120 | 4 | 53.11 | 63.25 | 21.3 |
| 22* | 52 | F | 130 | 6 | 47.57 | 43.27 | 19.0 |
| 23* | 24 | M | 150 | 3 | 55.45 | 59.56 | 21.9 |
| 24* | 38 | F | 150 | 2 | 55.61 | 66.67 | 34.9 |
| 25* | 25 | M | 140 | 6 | 51.51 | 44.35 | 22.9 |
| 26 | 25 | F | 150 | 5 | | | 27.8 |
| 27* | 41 | F | 90 | 1 | 57.01 | 68.53 | 28.3 |
| 28 | 27 | F | 70 | 4 | 51.76 | 44.46 | 28.2 |
| 29* | 26 | M | 140 | 3 | 58.15 | 62.48 | 28.2 |
| 30* | 76 | M | 90 | 5 | 61.02 | 67.91 | 25.8 |
| Mean | 33.3+/-14.8 | | 132+/-32.8 | 3.70+/-1.20 | 55.6+/- 3.50 | 61.2+/-6.70 | 23.7+/-2.20 |

**Table 2 | List of features input to XGBoost model.**

| Feature # | Type | Feature |
|---|---|---|
| 1 | Solo Features | Heart rate |
| 2 | | Amplitude |
| 3 | | Systolic Time |
| 4 | | Diastolic Time |
| 5 | | F3* F4 |
| 6 | | (F3/F4)*F1 |
| 7 | | Systolic upslope |
| 8 | | F3*F1 |
| 9 | | Time from systolic base to max systolic upslope |
| 10 | | Time from max systolic upslope to systolic peak |
| 11 | | F9/F10 |
| 12 | | FWHM at 50% of peak amplitude |
| 13 | | FWHM at 25% of peak amplitude |
| 14 | | FWHM at 75% of peak amplitude |
| 15 | | Systolic area |
| 16 | | Diastolic area |
| 17 | | Systolic/Diastolic area |
| 18 | | First harmonic amplitude |
| 19 | | Second harmonic amplitude |
| 20 | | Third harmonic amplitude |
| 21 | | Second harmonic phase- first harmonic phase |
| 22 | | Third harmonic phase- first harmonic phase |
| 23 | | Third harmonic phase- second harmonic phase |
| 24 | | F20/F18 |
| 25 | | F19/F18 |
| 26 | | F20/F19 |
| 27 | Joint features | Time delay between finger BFi and PPG |
| 28 | | Time delay between wrist and finger PPG |
| 29 | | Time delay between wrist BFi and finger BFi |
| 30 | | Time delay between wrist BFi and finger PPG |
| 31 | | Time delay between wrist BFi and wrist PPG |
| 32 | | Time delay between finger BFi and wrist PPG |
| 33 | | FH phase PPG finger - FH phase BFi finger |
| 34 | | FH phase PPG wrist- FH phase BFi wrist |
| 35 | | FH phase BFi finger - FH phase BFi wrist |
| 36 | | FH phase PPG wrist - FH phase BFi finger |
| 37 | | FH pahse BFi wrist - FH pahse PPG finger |
| 38 | | FH phase PPG wrist - FH phase PPG finger |
| 39 | | F4 BFi * F3 PPG from finger |
| 40 | | F3 BFi * F3 PPG from finger |

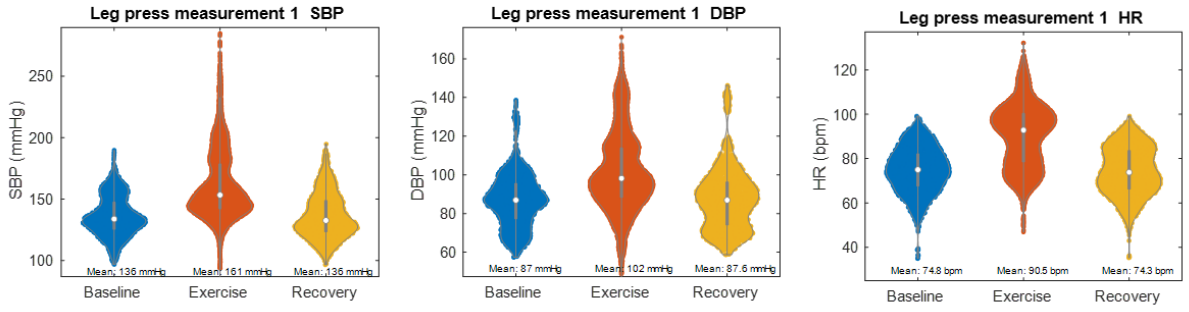


**Figure 1 | SBP, DBP, and HR distributions during initial leg press measurement.** Left to right: SBP, DBP and HR distributions during the first leg press measurement for N = 30 subjects. Mean values at each condition are shown on plots.

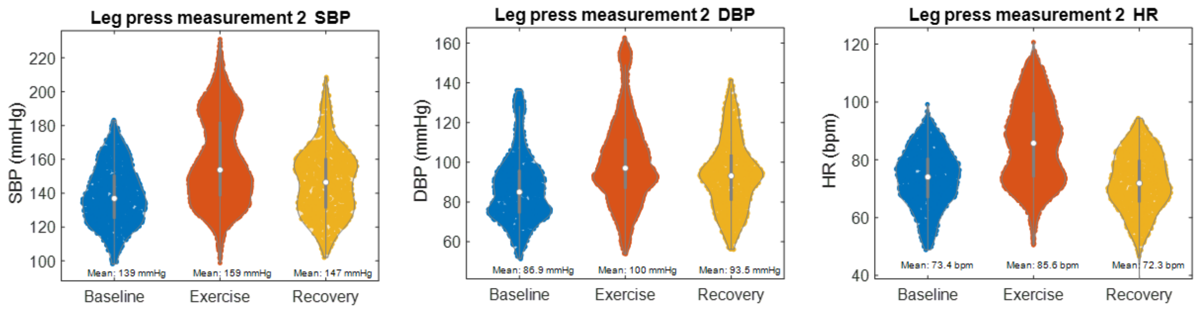


**Figure 2 | SBP, DBP, and HR distributions during second leg press measurement.** Left to right: SBP, DBP and HR distributions during the first leg press measurement for N = 20 subjects. Mean values at each condition are shown on plots.

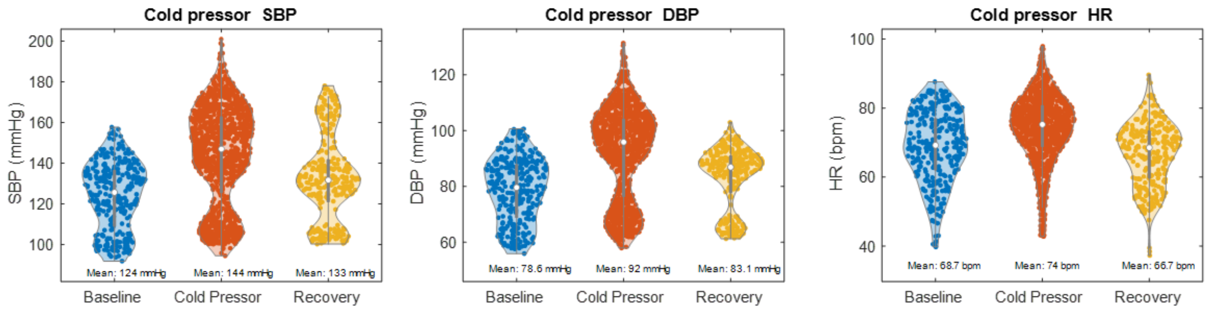


**Figure 3 | SBP, DBP, and HR distributions during cold pressor measurement.** Left to right: SBP, DBP and HR distributions during the first leg press measurement for N = 5 subjects. Mean values at each condition are shown on plots.

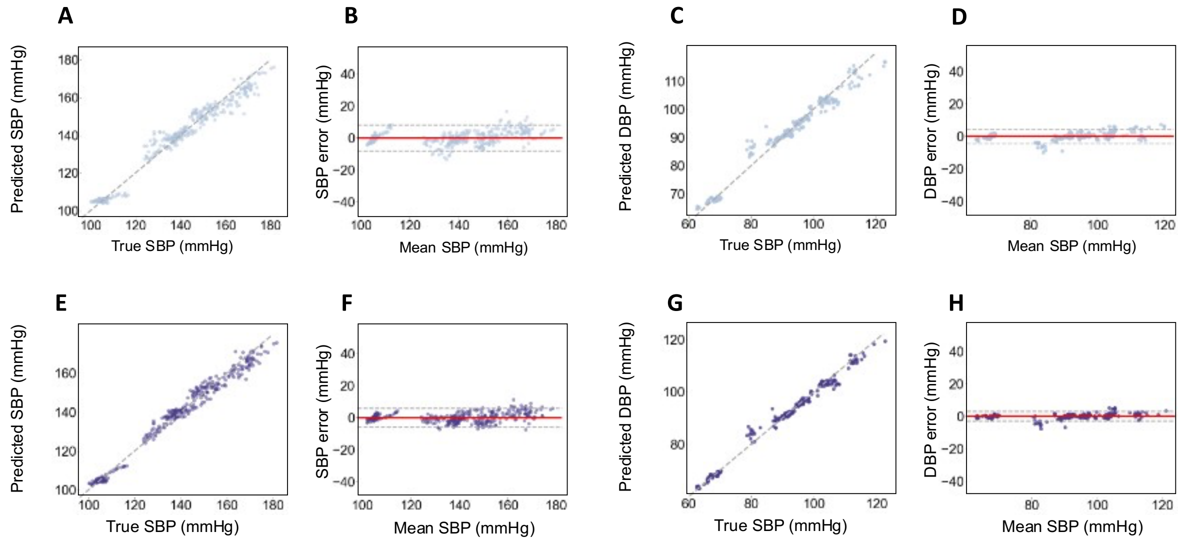


I

| | SBP | | | | DBP | | | |
|---|---|---|---|---|---|---|---|---|
| | Grand errors | | Subject errors | | Grand errors | | Subject errors | |
| | PPG | PPG + BFi | PPG | PPG+ BFi | PPG | PPG + BFi | PPG | PPG+ BFi |
| ME (mmHg) | 0.11 | 0.16 | 0.06 | 0.18 | 0.11 | 0.13 | 0.13 | 0.16 |
| SD (mmHg) | 4.18 | 3.02 | 4.23 | 3.01 | 2.55 | 2.43 | 2.61 | 2.55 |
| MAE (mmHg) | 3.16 | 2.33 | 3.57 | 2.46 | 2.01 | 1.93 | 2.14 | 2.01 |

**Figure 4 I Cold pressor SBP and DBP results. a.** Correlation between predicted and true SBP for the PPG model. **b.** SBP Bland – Altman plot for the PPG model. **c.** Correlation between predicted and true DBP for the PPG model. **d.** DBP Bland – Altman plot for the PPG model. **e.** Correlation between predicted and true SBP for the BFi + PPG model. **f.** SBP Bland – Altman plot for the BFi + PPG model. **g.** Correlation between predicted and true DBP for the BFi + PPG model. **h.** DBP Bland – Altman plot for the BFi + PPG model. **i.** Table with SBP and DBP grand and subject errors.

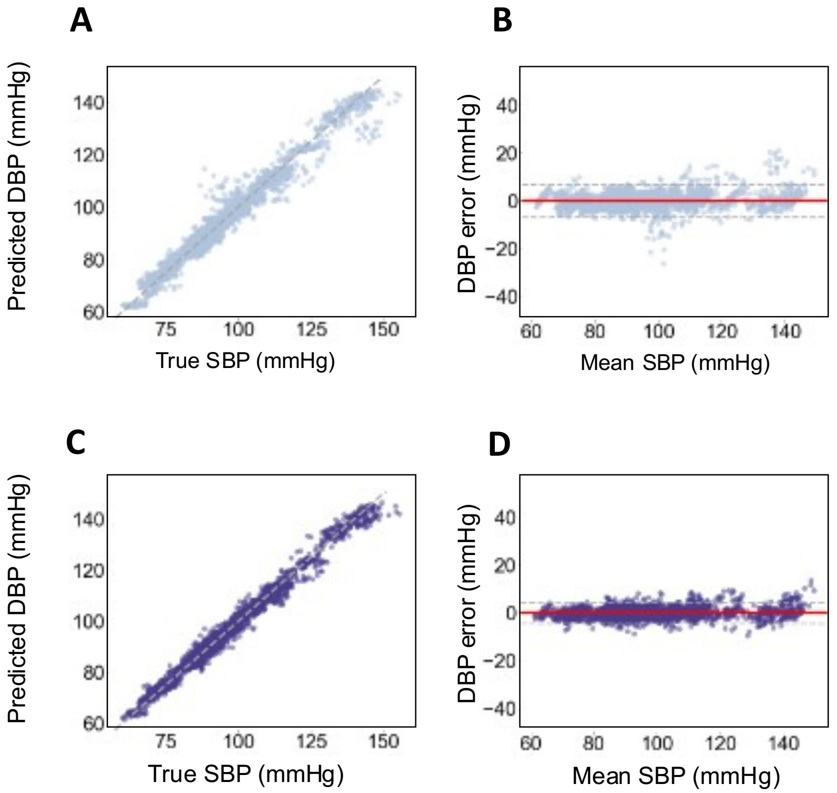


**E**

| | DBP | | | |
|---|---|---|---|---|
| | **Grand errors** | | **Subject errors** | |
| | PPG | PPG + BFi | PPG | PPG+ BFi |
| ME (mmHg) | 0.12 | 0.10 | 0.12 | 0.09 |
| SD (mmHg) | 3.46 | 2.27 | 2.96 | 2.14 |
| MAE (mmHg) | 2.40 | 1.69 | 2.45 | 1.69 |

**Figure 5 I 5-fold cross validation DBP results. a.** Correlation between predicted and true DBP for the PPG model. **b.** DBP Bland – Altman plot for the PPG model. **c.** Correlation between predicted and true DBP for the BFi + PPG model. **d.** DBP Bland – Altman plot for the BFi + PPG model. **e.** Table with DBP grand and subject errors.

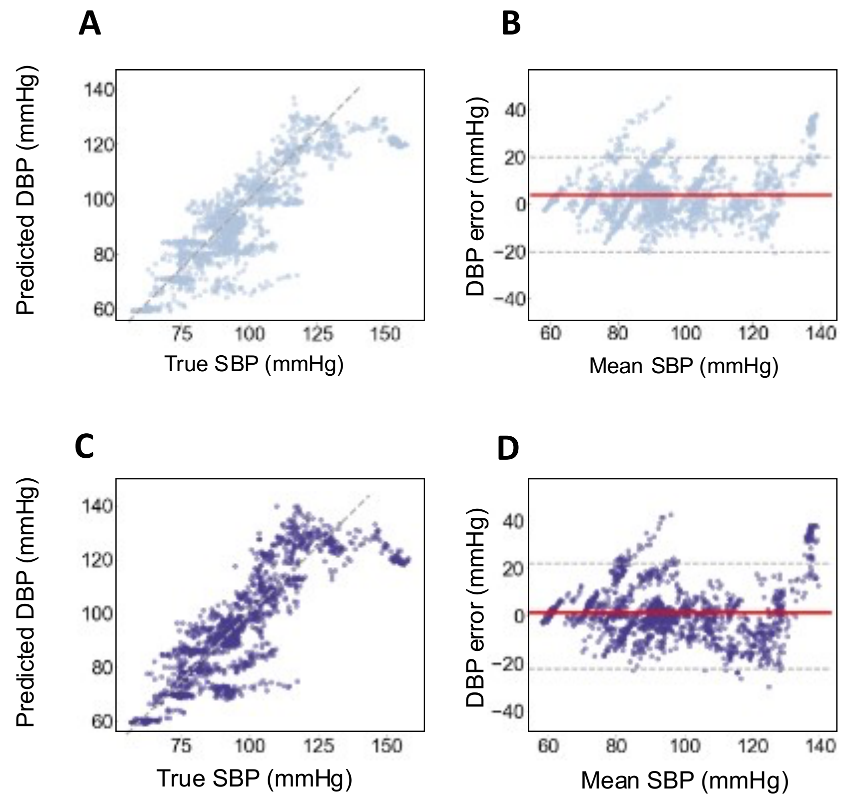


E

| | DBP | | | |
|---|---|---|---|---|
| | Grand errors | | Subject errors | |
| | PPG | PPG + BFi | PPG | PPG+ BFi |
| ME (mmHg) | 3.91 | 1.40 | 3.50 | 1.53 |
| SD (mmHg) | 10.3 | 11.28 | 6.15 | 6.62 |
| MAE (mmHg) | 7.77 | 8.00 | 7.64 | 7.68 |

**Figure 6 I Longitudinal DBP results. a.** Correlation between predicted and true DBP for the PPG model. **b.** DBP Bland – Altman plot for the PPG model. **c.** Correlation between predicted and true DBP for the BFi + PPG model. **d.** DBP Bland – Altman plot for the BFi + PPG model. **e.** Table with DBP grand and subject errors.

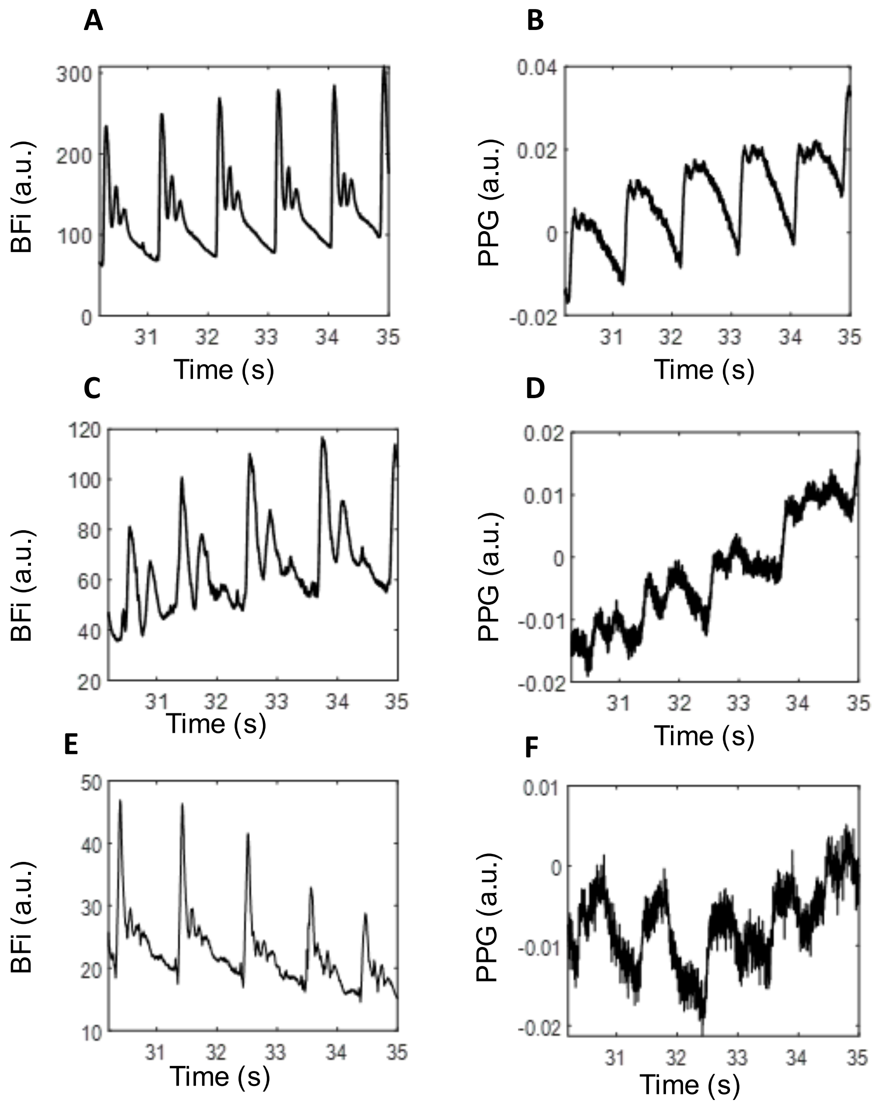


**Figure 7 | Finger BFi time traces from 3 subjects showing the differences in noise between BFi and PPG pulse waveforms. a.** Finger BFi from subject 2. b. Finger PPG from subject 2. c. Finger BFi from subject 17. d. Finger PPG from subject 17. e. Finger BFi from subject 19. f. Finger PPG from subject 19.

**Table 3 l Top 25 features sorted by sum of importances.** For each of the 20 remeasured subjects, 25 features are selected. We extracted importances for all of the features selected across the 20 remeasured subjects. The importances for each feature were then summed across each occurrence. For example, the top feature occurred was chosen for the model in 12 of the remeasured subjects (Frequency = 12). The importance for each of those occurrences was summed to get the sum of importances (Σ Importances = 1.02). The top 25 features ranked by importance sum are shown in the table below.

| | Feature | Type | Frequency | Σ Importances |
|---|---|---|---|---|
| **1** | Third harmonic phase - first harmonic phase | Finger BFi | 12 | 1.02 |
| **2** | Heart rate | Finger BFi | 6 | 1.01 |
| **3** | Diastolic Area | Wrist PPG | 8 | 0.87 |
| **4** | Second harmonic ratio | Finger BFi | 6 | 0.86 |
| **5** | Diastolic time * systolic time | Finger BFi | 7 | 0.52 |
| **6** | Third harmonic amplitude/Second harmonic amplitude | Finger BFi | 13 | 0.50 |
| **7** | Amplitude | Wrist BFi | 12 | 0.50 |
| **8** | Amplitude | Finger BFi | 10 | 0.48 |
| **9** | Width of pulse at 25% of amplitude | Finger BFi | 9 | 0.46 |
| **10** | Slope | Finger BFi | 12 | 0.41 |
| **11** | Width of pulse at 50% of amplitude | Finger BFi | 8 | 0.41 |
| **12** | Slope | Wrist BFi | 9 | 0.37 |
| **13** | Second harmonic ratio | Wrist BFi | 4 | 0.37 |
| **14** | Systolic/diastolic area | Finger BFi | 9 | 0.36 |
| **15** | Diastolic time * systolic time | Wrist PPG | 4 | 0.35 |
| **16** | Second harmonic amplitude | Wrist BFi | 4 | 0.35 |
| **17** | Third harmonic amplitude/Second harmonic amplitude | Wrist PPG | 8 | 0.34 |
| **18** | Amplitude | Wrist PPG | 6 | 0.34 |
| **19** | Heart rate | Wrist PPG | 5 | 0.33 |
| **20** | Third harmonic amplitude | Wrist PPG | 7 | 0.31 |
| **21** | Third harmonic ratio | Wrist PPG | 4 | 0.29 |
| **22** | Diastolic time | Finger BFi | 8 | 0.29 |
| **23** | Third harmonic amplitude/Second harmonic amplitude | Finger PPG | 3 | 0.27 |
| **24** | Diastolic Area | Finger BFi | 7 | 0.27 |
| **25** | Width of pulse at 25% of amplitude | Wrist BFi | 7 | 0.27 |

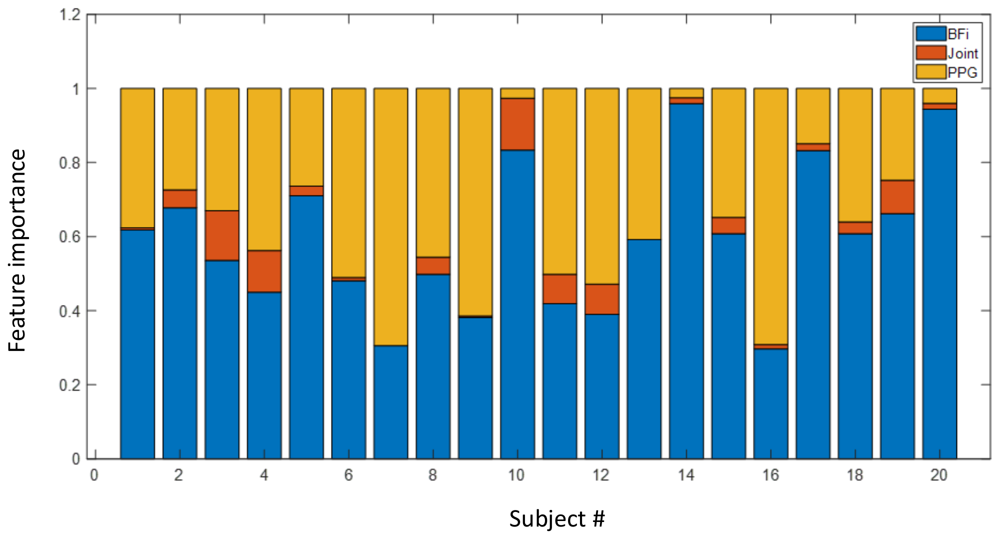


**Figure 8 l Sum of BFi, PPG and Joint feature importances for each of the 20 remeasured subjects.** For each of the 20 remeasured subjects, 25 features are selected. We extracted importances for all of the features selected across the 20 remeasured subjects. The importances of BFi features, PPG features, and Joint features were summed. BFi and Joint features have proportionately higher importance in the models for 15 of the 20 remeasured subjects.
